# Preliminary Minimum Reporting Requirements for Reporting In-Vivo Neural Interface Research: I. Implantable Neural Interfaces

**DOI:** 10.1101/2020.11.18.375741

**Authors:** Calvin D. Eiber, Jean Delbeke, Jorge Cardoso, Martijn de Neeling, Sam E. John, Chang Won Lee, Jerry Skefos, Argus Sun, Dimiter Prodanov, Zach McKinney

## Abstract

The pace of research and development in neuroscience, neurotechnology, and neurorehabilitation is rapidly accelerating, with the number of publications doubling every 4.2 years. Maintaining this progress requires technological standards and scientific reporting guidelines to provide frameworks for communication and interoperability. The present lack of such standards for neurotechnologies limits the transparency, reproducibility, and meta-analysis of this growing body of research, posing an ongoing barrier to research, clinical, and commercial objectives.

Continued neurotechnological innovation requires the development of some minimal standards to promote integration between this broad spectrum of technologies and therapies. To preserve design freedom and accelerate the translation of research into safe and effective technologies with maximal user benefit, such standards must be collaboratively co-developed by a full spectrum of neuroscience and neurotechnology stakeholders. This paper summarizes the preliminary recommendations of IEEE Working Group P2794, developing a Reporting Standard for *in-vivo* Neural Interface Research (RSNIR).

**Impact Statement:** This work provides a preliminary set of reporting guidelines for implantable neural interface research, developed by IEEE WG P2794 in open collaboration between a range of stakeholders to accelerate the research, development, and integration of innovative neurotechnologies.

## I Introduction

NEURAL interfaces (NIs) are systems that record and/or modulate the activity of the nervous system (see Fig. 1). A broad spectrum of technological modalities for NIs have been developed over the last 50 years, including both invasive (implanted) and non-invasive systems. NIs have been shown to provide therapeutic benefit for a wide range of conditions, as well as providing powerful tools for studying nervous system physiology, improving human-machine interaction, and augmenting human capabilities [1]. The rapid proliferation of neurotechnology in recent years (Fig. 1B) has produced a wealth of devices and systems with advanced neurosensing and neuromodulatory capacities, with a wide range of potential clinical and consumer applications. This diversity of NI technologies, applications, performance metrics, and experimental paradigms – along with the present lack of technological standards and reporting guidelines – has severely limited the transparency, reproducibility, and meta-analysis of this body of research and hampered its translation into widely beneficial and commercially available neurotechnologies.

**Fig. 1.**
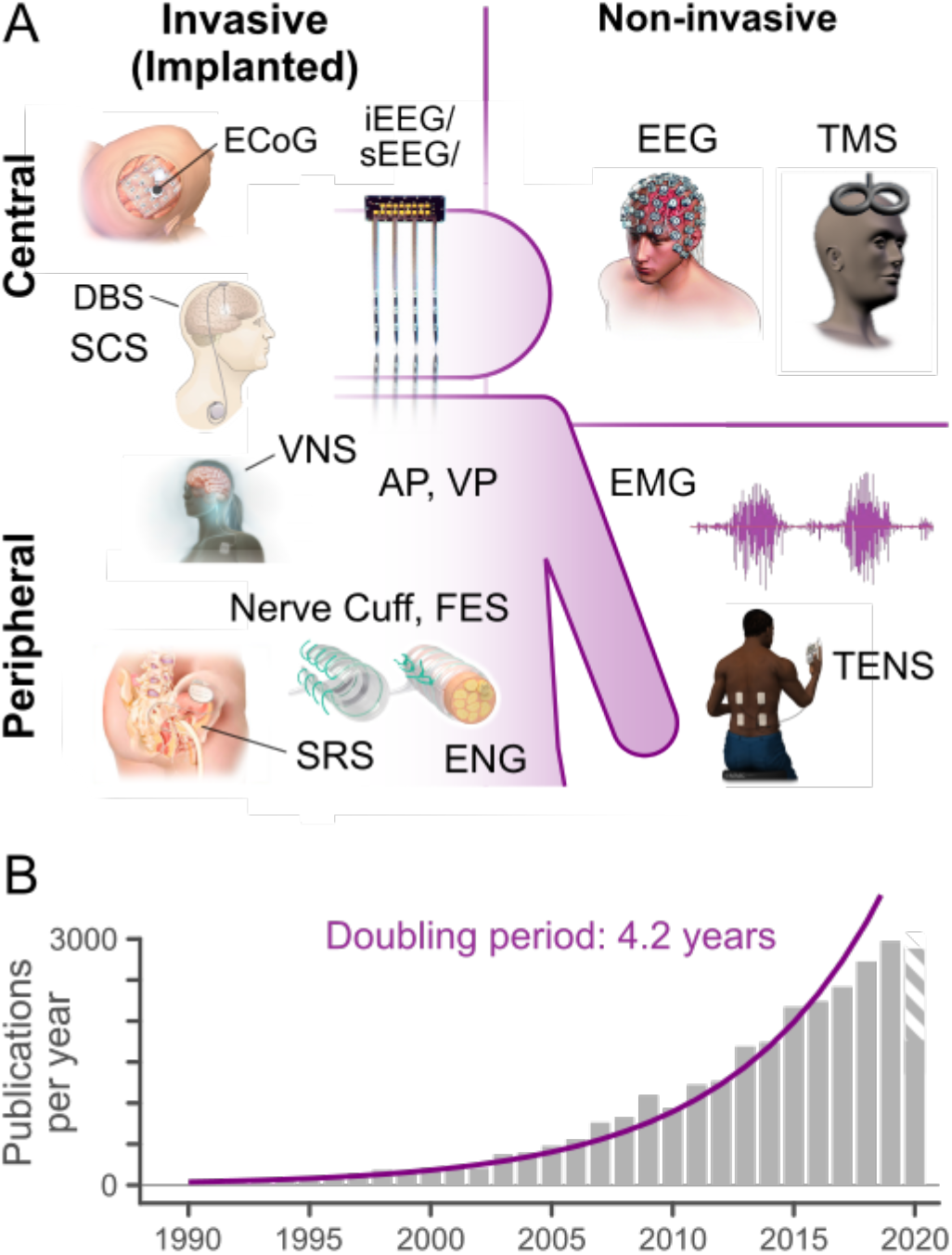
A) Overview of common NI technologies and applications. Neurosensing Modalities: EEG (electroencephalography), ECoG (electrocorticography), i/sEEG (intracranial/stereotaxic EEG), EMG (electromyography, ENG (electroneurography). Neuromodulation modalities: AP (auditory prosthesis), DBS (deep brain stimulation), FES (functional electrical stimulation), SCS (spinal cord stimulation), SRS (anterior sacral root stimulation), TENS (transcutaneous electrical nerve stimulation), TMS (transcranial magnetic stimulation), VNS (Vagus nerve stimulation), VP (visual prosthesis). B) The accelerating rate of growth for neural interface research (see supplemental methods), in publications per year.

The effective interpretation, aggregation, and metaanalysis of NI research thus requires more extensive reporting standards to improve the overall ‘information interoperability’ of NI study reports and data [2]. Many such reporting guidelines and initiatives have been enacted in recent years to address the so-called “replication crisis” across health and cognitive science research. For example, the Enhancing the Quality and Transparency of health Research (EQUATOR) network has compiled a list of best-practice reporting guidelines specific to different types of clinical and health-related studies, including the CONSORT guidelines for randomized clinical trials, the ARRIVE standard for pre-clinical animal trials, the PRISMA guidelines for systematic reviews, and many more. Regarding the sharing and interoperability of scientific data, the FAIR principles of findability, accessibility, interpretability, and reusability [3] have been widely endorsed [4] and represented in numerous neuroinformatics initiatives, including the International Neuroinformatics Coordinating Facility [5], Neurodata Without Borders, and the Brain Imaging Data Structure [6].

Despite this progress, the sum of existing standards and guidelines lacks the technical specificity to ensure a sufficiently detailed description of NI systems, methods, and results to ensure accurate interpretation and reproducibility. To address this ‘standardization gap,’ IEEE Standards Association Working Group (WG) P2794 – spawned from the IEEE Industry Connections Activity on Neurotechnology for Brain-Machine Interfaces [7] in parallel with WG-P2731 (Unified Terminology for Brain-Computer Interfaces) — is currently developing a set of Reporting Standards for *in vivo* Neural Interface Research (RSNIR), with the primary objective of improving the quality and transparency of NI research across a full spectrum of neurotechnological modalities. This standard is intended to establish the technological specificity necessary to achieve full interpretability and reproducibility of NI studies – and thereby to improve the scientific quality and impact of NI research in facilitating the development of safe and effective neurotechnologies.

While a primary application of this Standard will be to scientific publications, it is intended to serve as reference for any entity that seeks to improve the rigor and transparency of NI research, including regulatory bodies and funding agencies, as well as translation of NI research into medical devices. This report previews one such set of guidelines under development. Constructive feedback is welcomed from all neurotechnology stakeholders, including scientific, commercial, clinical, regulatory, and end-user perspectives.

## II. Scope

The official scope of the IEEE Working Group P2794 is to “define the essential characteristics and parameters of in-vivo neural interface research studies (including clinical trials) to be reported in scientific and clinical literature, including both minimum reporting standards and best-practice guidelines.” WG P2794 has defined the scope of NIs to be addressed by the Standard to include all engineered systems that directly record bio-signals of neurological origin and/or directly modulate neural activity. “NI research” is defined to include all studies where NI technologies are employed, either as the object of investigation or solely for recording data. More details regarding the scope and organization of P2794 are provided in the Supplementary Materials.

This article specifically sets forth a minimum information standard (in the FAIR [3] sense, e.g. [8], [9]) for reporting research involving implanted NIs. The technology underlying electrode-based NIs is more mature than other NI approaches [10], [11], so specific recommendations for reporting electrode-based NI research are provided. The scope of this module does not include aspects of NIs for which other standards have been provided, for instance in assessing biocompatibility [12] or characterization of research subjects [13]–[15].

## III. Reporting Topics for Implantable Neural Interfaces

To promote findable, accessible reporting [3], NI research publications should specify the NI technology(s), neuroanatomic targets, use paradigms /applications, and overall study design in the publicly-accessible metadata (title, abstract, and keywords). RSNIR-compliant NI study reports should adhere to all applicable reporting guidelines (e.g. EQUATOR [16], CONSORT [13], ARRIVE [14]). The purpose of the RSNIR standard is to expand upon these guidelines by identifying the technological and methodological details necessary to ensure clear, reproducible NI reporting. Accordingly, requirements already covered in these ‘parent’ guidelines will not be exhaustively listed here, but may be repeated for clarity and context.

### A. Neural Interfacing Context and Study Aims

To provide sufficient context and rationale, the background/introduction section of NI study reports should clearly identify the fundamental capabilities and limitations in the pertinent technological state-of-the-art and the scientific knowledge gaps addressed by the current study, with reference to authoritative works. Reports should specify the technological or methodological innovation(s) and scientific hypotheses proposed by the study. <u>Testable</u> hypotheses and additional qualitative/descriptive study aims should be stated in relation to the study’s primary outcome measures.

Along with aims, the developmental stage of the study (technology development [17] vs. animal studies [18] vs. clinical validation [15]) should be identified per Table 1. The report should indicate which NI application scenario(s) were investigated, per the IEEE NeuroEthics framework [19]:

- Recording/sensing (e.g. for scientific understanding or diagnosis),
- Stimulation/neuromodulation (e.g. to restore or enhance sensory, motor, or cognitive function)
- Closed-loop control of applications or prosthetic devices,
- Physical/biological modification
- Neural augmentation and facilitation.

**TABLE I.** Reporting Topics for NI Study Aims and Context

| Reporting Topic | High-level Descriptors | Detailed Descriptors |
| --- | --- | --- |
| Study Aims and Type | Foundational concept and technology development | Pre-clinical concept design study (e.g. human cadaver),<br>Modelling & simulation of NI performance,<br>Benchtop evaluation of NI capabilities and reliability. |
|  | Demonstration in animal models | Acute animal validation and refinement of mechanism,<br>Chronic passive safety and reliability,<br>Chronic active full system test (ideally in a disease model). |
|  | Validation and confirmation in clinical settings | Acute clinical safety, partial intra-operative test,<br>Feasibility & verification (pilot study),<br>Validation study (pivotal study / clinical trial),<br>Evaluation & monitoring (post-market) |
| Intended Application | Neuromodulation (stimulation) | Sensory neuromodulation (e.g. cochlear prosthesis),<br>Motor (Efferent) neuromodulation (e.g. FES). |
|  | Neurosensing (recording) | Diagnostic (e.g. epileptic foci discrimination),<br>Control of an external prosthesis<br>Control of virtual applications. |
|  | Closed-loop control or operation | Diagnostic (e.g. H-wave, epilepsy),<br>Targeted delivery of therapy<br>Sensorimotor integration |
| Physical Modality /<br>Technology | Electrical | quasi-electrostatic (μs-s timescales), tDCS<br>electrodynamic (fs-ns timescales, e.g. [35]),<br>single- or multi-unit recording,<br>field potential recording, ECoG, EEG. |
|  | Magnetic and Electromagnetic | fMRI, MEG, TMS |
|  | Optical and Infrared | Optogenetic stimulation<br>Voltage-sensitive or calcium-sensitive recording<br>fNIRs, IR stimulation |
|  | Acoustic | Focused ultrasound stimulation |
| Target Structure | Central Nervous System (CNS) | Targeted brain or spinal cord region(s) to be named per [27]-[29] |
|  | Peripheral Nervous System (PNS) | Targeted neuroanatomical structures to be named per [27]-[29] |

These loosely align with the application scenarios for braincomputer interfaces identified in [1]: replacing, restoring, enhancing, supplementing, improving, and studying neurological function. Finally, The introductory NI description should specify the target neuroanatomical structure(s) and device-tissue interface type/region.

### B. NI Experimental Design and Outcome Measures

As a guiding principle, all aspects of experimental designs featuring NIs should be described in sufficient detail to permit replication by other researchers. The number and type of subjects involved in the study must be clearly stated, along with the other characteristics listed in table 2. All NI studies must comply with consensus standards of ethical conduct, including local regulations, institutional review board approval, and the Declaration of Helsinki [20].

**TABLE II.** Reporting Topics for NI Experimental Design and Outcome Measures

| Reporting Topic | High-level Descriptors | Detailed Descriptors |
| --- | --- | --- |
| Human Subjects | Complete list of clinical criteria | Complete list of inclusion and exclusion criteria,<br>Criteria used to allocate subjects to experimental groups<br>Recruitment methods for subjects and controls |
|  | Demographic characteristics | Number and type of subjects involved,<br>Subject age and gender<br>Statistical justification of sample size, including “convenience sampling” |
|  | Relevant clinical history | Timelines of disease onset and symptom presentation<br>All administered drugs and drug doses, including administration routes.<br>Any other parallel treatments<sup>1</sup>. |
| Animal Models | Fundamental characteristics | Number and type of subjects involved, including justification of sample size.<br>Species/ strain, bodyweight, genetic manipulations (if relevant). |
|  | Husbandry and housing conditions | light/ dark schedule, environmental enrichment, experimental location.<br>All administered drugs and drug doses, including administration routes<sup>1</sup>. |
|  | Training and behavior (if relevant) | Training, reward, and performance assessment methods. |
| Interventions | Description of all interventions applied | (procedures, devices, treatment programs, surgical procedures , etc.) |
|  | Sequential timeline of interventions | including sequences and interrelations,<br>Randomization for cross-over / within-subjects-type designs |
|  | Location and setting of the experiments | (e.g. clinic, home setting, animal laboratory or home cage) |
|  | Experimental Equipment (Excluding NI) | Any specialized medical equipment used during the experiments,<br>Experimental stimulator / actuator and system information,<br>Any other sensors or actuators used to assess the performance of the NI: Vendor, make and model, control or acquisition system software and version |
| Stimulus Description | Visual Stimuli<sup>2</sup> | Background illumination level (e.g. scotopic or in units of cd / m2), Adaptation state of the experimental subject (e.g. dark-adapted),<br>Duration of the stimulus including any adaptation or masking procedures,<br>Approximate retinotopic location of the stimulus presentation (e.g. foveal)<br>Frame-rate and luminance range of the display. |
|  | Auditory Stimuli<sup>2</sup> | Background and stimulus sound levels,<br>Stimulus presentation (e.g. monaural, binaural),<br>Tone frequency and duration. |
|  | Tactile Stimuli | Similarly, for tactile stimuli, the stimulus type (e.g. vibratory, single-pulse, von Frey, etc.), intensity (in mm/s) and other properties should be reported. |
|  | Other Stimuli | For more complex stimuli, such as movies or sequences of spoken words, examples should be provided as supplementary data. |
| Outcome Measures | Basic signal quality metrics for NIs | (e.g. signal-to-noise ratio) |
|  | Usability and patient satisfaction scores. | For animals research, these may include behavioral assessments e.g. [36] |
|  | Computation of derived measures | References to established measures and formulas for novel measures |
| Statistics | Identification of dataset(s) between which each comparison was conducted | Description and rationale for data grouping provided (e.g. between vs. within-subjects comparisons). |
|  | Derivation for each datum | Time point(s) for data sampling<br>Single measurement or aggregated measures. |
|  | Other statistical methods | Methods used to examine subgroups,<br>Assessment of multi-variate interactions,<br>Control for confounding and missing data,<br>Mitigation of potential sources of bias. |

<sup>1</sup> This is important, as many drugs have effects on the nervous system which may influence NI performance, e.g. [37]
<sup>2</sup> For more complex stimuli, such as movies or sequences of spoken words, examples should be provided as supplementary data.

#### B.1. High-level study design

The NI study description should first identify the overall experimental design type(s) using established paradigms such as single/double-cohort, crossover, withdrawal or longitudinal study designs [21]. Within-subjects designs (where each participant serves as their own control, such as n-of-1 case studies [22]) are common for early clinical and pre-clinical NI research; the main motivation being to demonstrate proof-of-concept and/or subject specific safety and effectiveness of the NI prior to conducting a large-scale clinical trials. Given the high tendency for individual variability, this approach demands a detailed description of the clinical and demographic characteristics of all subjects. Follow-up data collection to monitor the clinical evolution after experimental intervention is highly encouraged (e.g. [23]).

Later-stage, larger-scale clinical studies intended to evaluate an intervention’s efficacy with respect to an established standard therapy for a broader user population typically employ a between-subjects study design, such as the “gold standard” randomized-controlled trial. Important for these types of experiments is the definition and recruitment of a representative control group. The use of placebo groups and blinded assessment of outcomes is strongly encouraged. This type of experiment can also be used in animal studies. In “crossover” designs featuring multiple interventions administered in serial, randomization of intervention sequence between subjects is advised, with a sufficiently long “washout” period to combat carryover effects (such as improved performance due to longer exposure to the NI). Baseline outcome measures should be noted before the start of intervention.

#### B.2. Description of Intervention(S)

All interventions, including procedures, NI devices, treatment programs, and surgical procedures, must be described in detail to ensure reproducibility. Stimulation and recording protocols, including the conditions under which the experiment was conducted, must be reported. If visual, auditory, tactile, or other sensory stimuli were used in either experimental or control conditions, these stimuli must be described per table 2. Whenever the experimental design involves behavioral assessments, potential behavioral biases and mitigation strategies (whenever applicable) should be reported (eg. human handedness, education, expectations about the study).

#### B.4. Outcome Measures and Statistical Analysis

All outcome and performance assessment measures – both NI-derived and otherwise – must be precisely defined. The selection and relevance of all such measures to the study aims and hypotheses should be justified. Basic signal quality metrics for NI data (e.g, signal to noise ratio) are recommended, as are usability and patient satisfaction scores.

All statistical analyses conducted should be reported in accordance with pertinent high-level reporting guidelines (EQUATOR, etc.). Reporting of data-processing and statistical methods must be sufficient to reproduce the presented results from raw data. The data set(s) between which each statistical comparison was conducted (e.g. between vs. within-subjects comparisons) must be clearly reported and justified. Where feasible, intended analyses of outcome measures should be documented and disclosed in advance of data collection in order to maximize transparency and the statistical validity of the results obtained and minimize the opportunity for so-called ‘p-hacking’.

### C. Description of the Neural Interface

The interpretability and reproducibility of NI research depends on accurate and complete descriptions of the NI in question. Underreporting of the device characteristics, particularly for clinical research, is the biggest barrier to reproducibility and meta-analyzability to NI research. To overcome this barrier, researchers must provide a thorough description of the NI (see table 3), including the specifics of the applied stimulus or recording procedures. These parameters are critical to comparing NI performance across technologies, devices, and cohorts.

**TABLE III.**
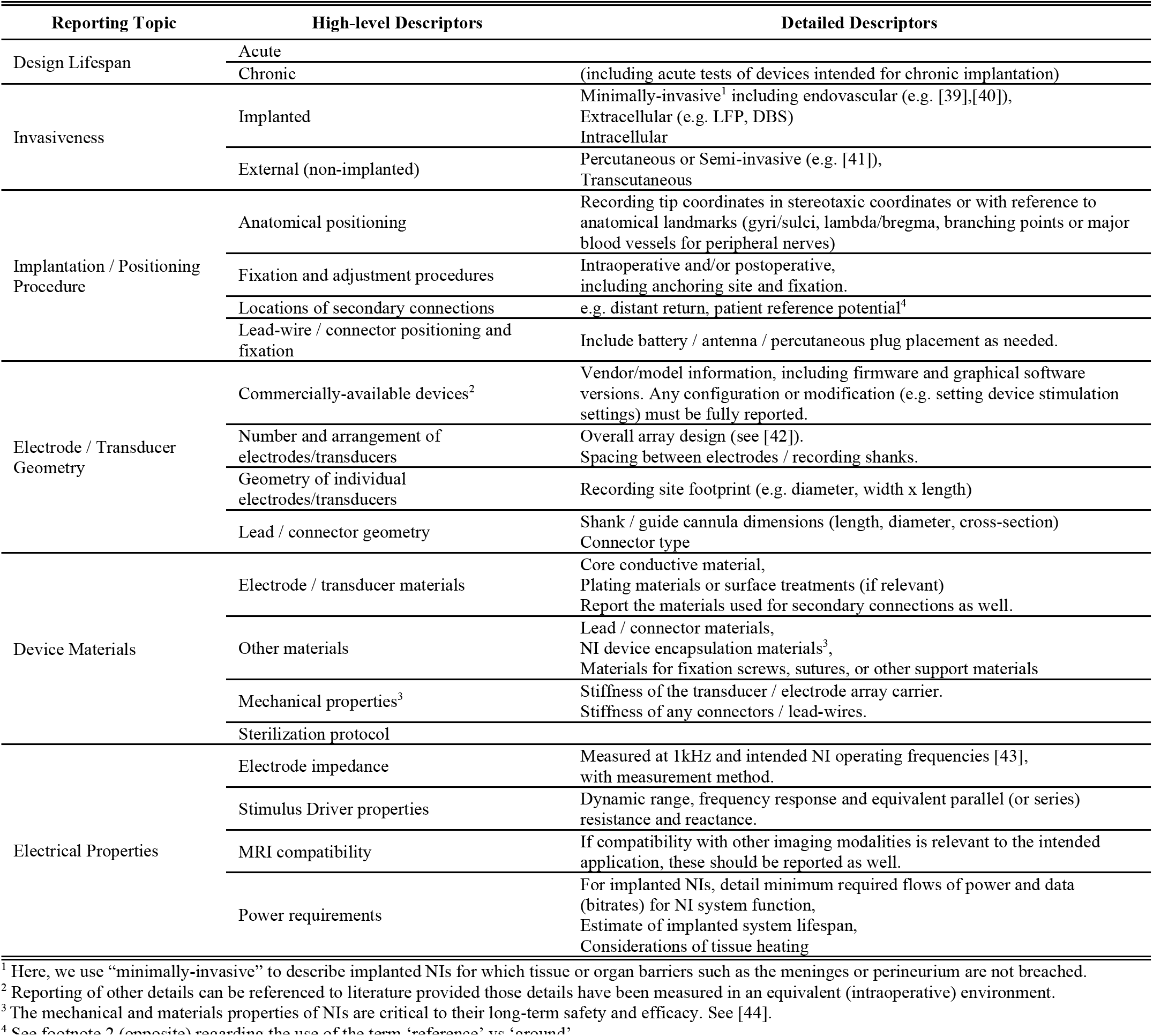
Reporting Topics for NI Physical Device Properties

Figure 2 shows a block diagram of a generic closed-loop NI system architecture which includes transducers (electrodes), signal acquisition and processing for neural recording, and stimulus generation and delivery for neuromodulation. The characteristics of all of these modules are essential for interpreting NI performance; essential reporting parameters for NI transducers and hardware is given in table 3, and essential reporting parameters for NI signal acquisition and processing is given in table 4. Diagrams such as figure 2 are essential for communicating the overall plan for a given NI approach and application, and we encourage their use for describing both the NI under test and the experimental context in which the NI is deployed. For custom experimental devices (including modified devices), authors should also provide a labelled diagram showing electrode / transducer sizes and locations^1^.

**Fig. 2.**
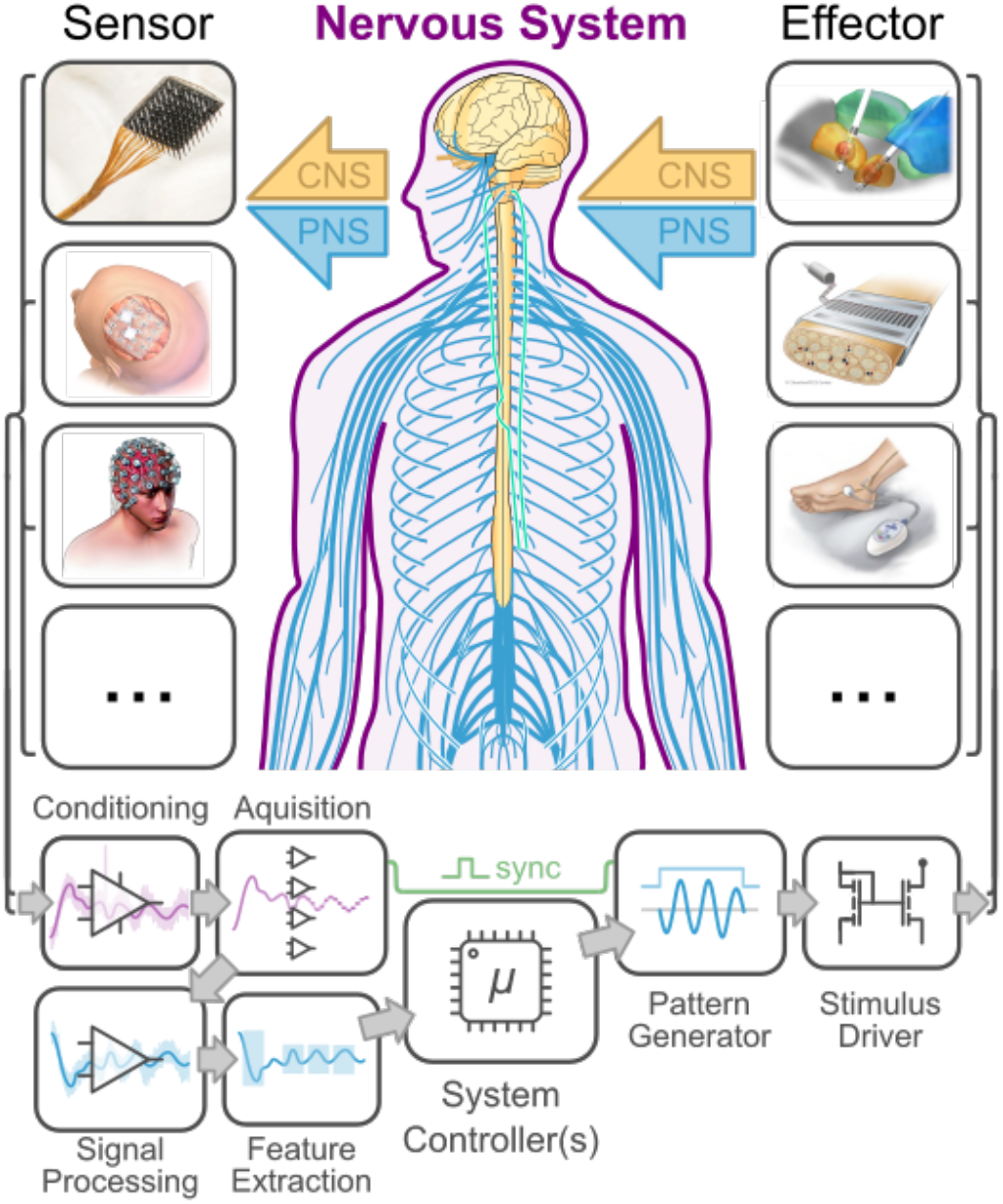
Block diagram of a prototypical NI system architecture. Sensors and effectors may interface invasicely or non-invasively with the central or peripheral nervous system (CNS / PNS). Neural sensing components will almost always include hardware signal conditioning, digital-to-analog conversion, digital signal processing, and feature extraction. Neuromodulation components include waveform selection and generation and the output drive to the stimulus end effector. Sensors, from top to bottom: high-density intracortical (Utah) array, ECoG array, EEG. Effectors: deep brain stimulation, peripheral nerve array (FINE, [38]), and transcutaneous stimulation.

**TABLE IV.** Reporting Topics for NI Signal Processing Properties

| Reporting Topic | High-level Descriptors | Detailed Descriptors |
| --- | --- | --- |
| Target Physiological Signal | High-resolution recording (e.g. single-unit) | Stability over time,<br>Cell type specificity and bias in recording. |
|  | Population-averaged recording (e.g. LFP, EEG, ECoG, ENG) | Stability over time,<br>Spatial and temporal resolution of the observed signal,<br>Cellular origin of the observed signal (e.g. [45]). |
|  | Neuromodulation | Spatial resolution of the imposed signal,<br>Cell type specificity of the evoked response,<br>Safety limits (power and/or current density [46]) and dynamic range,<br>Observed threshold ranges.<br>For nerve block, what is the purported mechanism of blockade? |
| Hardware Conditioning and Acquisition | Filtering | Input gain, anti-aliasing filter time-constant<sup>1</sup><br>Hardware artifact rejection and saturation range<sup>2</sup> |
| | Analog to digital conversion | Sampling rate, dynamic range, resolution (in bits or $\mu V$) |
|  | Output channels | Number of channels,<br>Map from sensor signals to channels (electrode/transducer identifiers)<br>Output channel characteristics including estimates of cross-tal |
| Signal Processing | Filtering | Filter type (high-pass, low-pass, band-pass, notch)<br>Details of any applied averaging or normalization<br>Details of any nonlinear filtering e.g. Kalmann filtering.<br>Visualize input and output signal characteristics |
|  | Artifact removal | Algorithm and parameter estimation<br>Order of applied signal-processing steps* |
| Feature Extraction | Frequency-domain transformations | Define all analysis bands used<br>Define wavelets and analysis windows for time-resolved spectral power estimation (e.g. [47]) |
|  | Spatial transformation | (e.g. inverse source localization) |
|  | Coordinate transformation | (e.g. PCA/ICA, SVD [48]).<br>Input and output signal characteristics |
|  | Datatype conversion | (e.g. spike detection and sorting, autoregressive model fitting)<br>Input and output signal characteristics<br>Sensitivity to input noise, signal-to-noise ratio. |
| Classification and Decision-making | Classification analyses | Classifier architecture (e.g. SVM, K-means, CNN, etc. see [50]).<br>Model parameters & hyperparameters<br>Initialization, Convergence criteria |
|  | Inference of mental states | The states identified must be clearly specified.<br>The usual 'expert' rules applied for offline analysis should be referenced (e.g. [49]) |
|  | Control Algorithms for effectors / stimulus delivery | Including algorithms for closed-loop control, "model-in-loop" control. |
| Stimulus Waveform Generation | Timing of stimulus delivery | Patient / subject control,<br>Trigger signals or events<br>Synchronization with recording equipment |
|  | Stimulus type | (e.g. current, voltage, optical, acoustic, etc.).<br>For electrical stimuli, equivalent standard rectangular current-controlled stimulus. |
|  | Timing of individual stimuli | Phase width, pulse shape, leading phase, inter-phase gaps.<br>Stimulus frequency, number of discrete stimuli per stimulation period<br>Shorting / charge-balancing procedures for electrical stimuli. |
|  | Output channels | Number of independently addressable stimulation channels available,<br>Electrode/transducer configurations used. |

<sup>1</sup> For recording physiological signals, transient artifacts can mimic physiological signals when filtered through high-pass filters higher than first-order. To avoid this, characterizing any such filter by a single time constant value can ensure this good practice has been enforced.
<sup>2</sup> If recording signals reach saturation during regular use (e.g. due to stimulation artifacts), this should be noted along with the expected duration of invalid signal.
<sup>3</sup> In principle, the order of linear signal processing steps is not important. In practice, malfunction, artefacts, and other sources of confusion are more easily identified in the frame of an orderly description.

The placement and positioning of the NI are critical to NI effectiveness (see [24]–[26]) and must be carefully reported (including the transducer, connectors, and any implanted electronics). Anatomical structures should be specified with reference to a widely-accepted formal vocabulary such as FIPAT [27] or recognized anatomical atlases (e.g. [28], [29]). Implantation and device positioning procedures must be described, including the location of each component relative to anatomical landmarks, expected error margins, and any criterion for surgical re-positioning or exclusion. Describe any procedures carried-out to confirm device position during or after concluding the experiment (e.g. histology, CT imaging). Finally, for research concerning entire implanted NI systems (as opposed to investigations of NI components), expected and observed implant lifespans should be reported, as well as any observed or predicted failure modes (e.g. [30]).

From a clinician, end-user, or regulatory perspective, the algorithms used for signal-processing, stimulus generation and closed-loop control are as much a part of a NI as the underlying hardware. Reporting of these aspects of NI systems must be conducted to the same level of rigor as reporting of the physical interface; essential reporting parameters are given in table 4. For neuro-sensing NIs, an unambiguous description of how signals from the electrodes / transducers are processed into recording channels is necessary. For novel NIs using recording approaches which might not be familiar to the wider NI community, the biophysical basis for the observed signals and measurement approach should be justified. Similarly, for novel neuromodulation NI approaches, the mechanism of the modulation of nerve activity should be described.

Algorithms used for signal conditioning, pre-processing, and analysis must be clearly reported and referenced. Links to public repositories containing open-source implementations with representative data sets are ideal. Inputs and outputs should be clearly specified, including confidence interval estimates (e.g. via bootstrap analysis of noisy input data, [31]). Existing standards for signal-processing research (e.g. [32]) should be applied.

### D. Neural Interface Results and Discussion

NI research reports should clearly and succinctly present the results of all analyses described in the methods (including primary and secondary outcomes), plus any additional post-hoc analyses (identified as such), in a manner that accurately summarizes and represents the full data set(s) analyzed, according to established biostatistical best practices [33], [34]. Graphical data representation (figures and tables) is preferred to text. Numerical values displayed in figures should be incorporated in the figure, a corresponding table, or supplemental materials. Wherever applicable (including aggregated measures and descriptive statistics), measurement variability and uncertainty should be quantified with standard measures (standard deviations, confidence intervals, etc.). Likewise, all comparisons conducting using inferential statistics should report statistical significance (or nonsignificance) and effect size. Where parametric statistics are used, the normality of data distribution should be confirmed. Rationale should be provided for the exclusion from presented analysis of any data collected within the same protocol. Measures of NI signal quality (e.g. signal-to-noise ratio) or essential performance are *strongly* recommended, along with presentation of example raw data.

All unexpected or adverse events (e.g. device failures or explantations, subject withdrawal, unplanned animal deaths, etc.) should be reported. Observed technical issues and complications should also be reported, including all mechanical, electrical, or software failures (broken electrodes, connections, etc.).

Discussion of results should address:

- To what extent do the results confirm the study hypothesis/es, and how do they fulfill the study objectives?
- Distinction between statistical and clinical (practical) significance, with reference to the observed effect size and uncertainty
- The fundamental novelty and/or significance of the findings with respect to the current state of the art, scientific body of knowledge, and/or field of potential applications. Comparisons to results of previous similar studies are encouraged, with attribution of notable similarities differences.
- The applicability and generalizability results to the intended NI users and applications, addressing concepts of validity (internal vs. external; construct; content; face)
- Discarded data collected according to the study protocol but excluded the final presented results/analysis
- Identification of key study limitations pertaining to the subject population, animal model, and/or experimental paradigm
  ○ Uncontrolled and potentially confounding factors
  ○ Precision and uncertainty of measurements, including intra-and inter-subject variability
  ○ The stability of neural recordings and/or stimulation parameters over the time course of the study
  ○ Potential sources of biases in the subject recruitment /enrollment process.
  ○ Study withdrawal rates
- Limitations of the presented technology/approach with respect to present or future application(s).
- Key challenges to the future development and application of the presented technologies, including usability considerations and open questions for further investigation.

## IV. Discussion

As a preview of IEEE Standard P2794 (RSNIR), this document has outlined minimum reporting requirements to ensure adequate transparency and reproducibility of *in vivo* research involving implantable neural interfaces (iNI). In this way, RSNIR complements existing scientific and clinical reporting guidelines by adding a layer of specificity to iNI technology. A majority of these recommendations may apply generically to all NI technology (including non-invasive modalities), and the RSNIR WG is currently working to adapt these requirements to such technologies, including EEG-based BCIs. In addition to scientific reporting guidelines, the RSNIR Standard will be supported by a network of complementary NI-relevant Standards under current development, including IEEE P2731 (Unified Technology for Brain-Computer Interfaces) and P2792 (Therapeutic Electrical Stimulation Waveforms). For medical NI technologies, RSNIR also aims to facilitate compliance with foundational medical device standards such as ISO 14971 (risk management), ISO 13485 (quality management systems), and IEC 60601 (safety and essential performance requirements).

The impact of RSNIR in promoting high-quality neuroscience and neurotechnology development depends critically on its widespread adoption by a range of institutions that define incentives across academic, commercial, and clinical domains, including high-impact scientific publications, funding agencies, regulatory bodies, and/or medical payers. To promote such adoption, the Standard seeks to define requirements to support an ‘ecosystem of information interoperability’ that serves the needs and objectives of all neurotechnology stakeholders, including aforementioned institutions as well as researchers, developers, clinicians, and end users.

To facilitate adoption at different levels of technological maturity (e.g. Technology Readiness Level [2]), RSNIR will apply the principle of *indirect reporting*, whereby reporting requirements may be fulfilled via reference to previous publications or documentation, provided that all required details are contained in the primary publication (including supplemental materials) and all others *directly* cited therein.

Regarding potential adoption by commercial entities, the RSNIR standard will seek to honor the proprietary nature of some NI system design details, by allowing the study reproducibility criterion to be fulfilled on a system-dependent basis – that is, by requiring the acquisition of commercial hardware or software. In such cases, public assurance of the NI system’s basic safety and performance may be achieved via third-party certification according to official testing Standards (UL, ASTM, etc.).

To make RSNIR usable and useful at all stages of research & development (technological maturity), feedback to this article and participation in the RSNIR WG are welcomed from all such stakeholders.

## Acknowledgments

We acknowledge Ricardo Chavarriaga, Jean-Louis Divoux, Rodolfo Fiorini, and all the other present and former members of WG 2794 who contributed in various ways to gathering the information condensed in this manuscript. We also acknowledge the administrative support from Tom Thompson (IEEE Standards Association) and Carole Carey (IEEE Engineering in Medicine & Biology Society).

## Footnotes

1 Electrode names like ‘anode’ and ‘cathode’ lead to confusion in the context of biphasic charge-balanced electrical stimulation to and should be avoided. Similarly, the labels ‘active’ or ‘reference’ imply assumptions about where activity or activation is occurring which may not be satisfied. For current-controlled stimuli, the term ‘return’ is clearer and should be preferred. For recording, ‘reference’ is to be preferred to other terms as this is the potential connected to a galvanically isolated recording device. As patients must never be connected to earth, by any means, terms such as ‘earth’, ‘neutral’, ‘safety ground’, or ‘building ground’ must not be used. We suggest using electrode / transducer labels like E1, E2, E3, etc.

