## Supplementary Materials for "Preliminary Minimum Reporting Requirements for Reporting In-Vivo Neural Interface Research: I. Implantable Neural Interfaces"

#### V.IEEE WG P2794 Scope and Organization

The RSNIR Working Group (WG) launched in January 2019 and remains active in the formulation of the proposed standards. The official scope of the WG is to "define the essential characteristics and parameters of in-vivo neural interface research studies (including clinical trials) to be reported in scientific and clinical literature, including both minimum reporting standards and best-practice guidelines." For the purposes of this standard, the RSNIR group has defined NI technologies as all engineered systems that record or modulate the activity of neural tissues, including the reading of biosignals of neurological origin (e.g. via electromyography, EMG).

The primary purpose of this Standard is to improve the interpretability, reproducibility, and meta-analysis of publicly reported in vivo research studies involving neural interfacing technologies (including both stimulation and recording from the central and peripheral nervous system) across different projects and institutions. Secondarily, this Standard aims to facilitate the further convergence and standardization of experimental methods, "benchmark" performance measures, and neurodata file formats, by defining the essential features of neural interface studies sufficient to render the research fully intelligible, reproducible, and conducive to further research.

To date, the group has been divided into 6 working sub-groups, comprising five oriented around the neurotechnological domains of EEG-based BCIs, implantable NIs, peripheral NIs, neuromodulation, and neuroimaging – plus a sixth group tasked with the 'horizontal integration' between the 5 technological 'verticals'. Topics to report include the design, configuration, and essential performance parameters of the neurotechnology(s) employed in the study, in addition to a thorough characterization of the experimental methodology, signal processing, and data analysis techniques. Notably, this Standard is intended to apply to research involving the use of NIs in any application, including both eliciting and measuring physiological responses and properties.

As a first step towards the definition of a reporting standard, the RSNIR WG has produced and internally reviewed a preliminary list of topics to be addressed by the Standard, intended to serve as the minimum reporting requirements necessary to render any NI study fully reproducible. At present, we are currently seeking feedback and refinement of this list from the neuroscientific and neurotech communities, via a complementary questionnaire. This survey is a putative list of topics which need to be included to ensure the interpretability and reliability of any report regarding a peripheral nerve interface, analogous to the reporting standards utilized within the EQUATOR network.

#### VI. Regarding Electrodes

**Electrode**. *n.* electricity conducting material in contact with biological tissue and connected through a lead to electrical equipment. In order to limit the electrical contact to a specific surface, electrodes are mounted in a probe made of insulating material.

Electrodes are used in two broad application categories: stimulation and recording. Although there is some overlap, the technical requirements usually differ between these cases.

##### *A. Stimulation (neuromodulation) electrodes*

The purpose of stimulation electrodes is typically to activate peripheral nerve fibers by depolarizing their axonal membranes. Except for the case of 'giant' axons and intra-cellular work, no membrane current can be applied directly. However, the depolarization (inward) current has been shown to be proportional to the second derivative of the electric field along the nerve fibers. Peripheral nerve stimulation is thus obtained by sending a current pulse through the surrounding volume conductor along the nerve. The stimulating inward membrane current is maximal under the negative electrode (often referred to as the cathode). Near threshold, the nerve will thus be activated under the negative electrode considered as the 'active' electrode while the second positive pole is considered as a 'reference'.

Because passing a current through an electrode in contact with a hydrous solution inevitably induces corrosion, stimulation pulses are typically biphasic with a second charge compensating pulse intended to 'undo' electrochemical changes in a period where the nerve activity has been triggered and is no longer sensitive to new stimuli. As a rule, a stimulus is thus biphasic, with two phases of opposite polarity. Note that in some conditions, under the anode, a membrane outward current can hyperpolarize an excited axon and thus block the further propagation of the generated action potential. Alternatively, high frequency currents can also induce such a propagation block, again under certain conditions.

The electrical field distribution in a volume conductor, especially when modified by the presence of the insulating components of a probe, can result in a virtual cathode, a spot where a membrane inward current is generated at distance from the real cathode.

*B. Recording (sensing) electrodes*

Action potentials correspond to short current pulses across the axon membrane. In the absence of a direct access to the inside of the membrane, the tiny electric field generated in the surrounding volume conductor can be recorded. These are so small, however, that recording is often at the limits of technical possibilities considering unavoidable noise and interferences. The recording instrumentation is differential, meaning it is arranged to see only the difference between two points at opposite sides of the source of interest but similarly exposed to external influences (the common mode). The result is that specifications such as noise figures, CMRR (common mode rejection ratio), and electrode impedance become critical. The common mode can be further reduced using a ground electrode connecting the base potential level of a floating pre-amplification stage at the level of the patient. It is important to note that recording quality and patient safety have a common requirement here in a good galvanic insulation of the patient-and-recording-input from the earth lead of the mains. These considerations are of course less important for implanted devices located inside the subjects.

Instead of a passive conductor, some devices replace the passive 'patient-ground' connection by an active system whereby an electrode can sense the potential level of the subject and use an active output to force that level to be identical to the recording input reference. This was first applied for ECG where sensing was obtained from an average of the electrode inputs and the active control feedback was place on the right leg, hence the name DRL (Driven right leg) now also applied in EEG practice.

*C. On electrode labels*

The discussion above should make it clear that electrodes should be named clearly with simple names that cannot substitute for a functional description and allowing to specify many electrodes. A simple number could do but an additional letter, for example S1, S2, S3.. for stimulation electrodes and R1, R2, R3..for recording electrodes allows to distinguish electrode names from a simple numbered list. G should be reserved for the 'patient ground' connection; calling this connection 'reference' creates confusion. An active electrode system needs a specific description of the sense input and the driven output.

*D. Montages and electrode function*

As indicated above, stimulation involves injecting a current into the body and signals are being recorded between at least two points. In each case, a single stimulation channel or a single recording channel involves two or more electrodes. In most instances one electrode corresponds to the procedure target, being either the point to be stimulated or the location of a physiological source to be recorded from. This is not always the case and sources can be considered as dipoles optimally located between the two electrodes. In all instances, in order to allow identification of the signal polarity, one electrode must be considered as target or active and the other as a reference (not to be confused with the 'patient ground'). It is important to realize that this is completely arbitrary but necessary to avoid confusion. All stimulation pulse and signal polarities should be defined according to the active or target electrode. In EEG, the arrangement of the electrodes of all channels is called a montage. That name can be extended to the peripheral situation. In brain applications, a single common reference is often used for all channels. This is not often the case in peripheral work but such arrangements as tripolar montages associate two electrodes connected together to the 'reference' input. These considerations pertain to the hardware arrangements. Software wise, simple subtractions between recorded channels allow to obtain any desired montage.

*E. Electrode localization*

Stimulation as well as recording involves a 3D potential distribution problem. Accurate electrode localization referring to the anatomical geometry is therefore essential. Standard electrode positions derived from measurements to anatomical landmarks are available for EEG (the 10/20 and 10/10 systems). No such generally accepted standard system is yet available for peripheral nerves or muscles, and precise localization is necessary to insure reproducibility. EMG textbooks describe optimal positions for muscle recordings, typically located over the end-plate. The muscle tendon is considered to offer an inactive spot. Other considerations must sometimes be taken into account such as the reduction cross-talk between muscles when a selective muscle is targeted or muscle activity considered as "noise" in a neural recording.

In addition to their anatomical localization, multi-electrode probes (arrays, multi-contact needles etc..) need a clear specification of the relative contact or electrode positions. Either a center-to center or an inter-contact (margin-to-margin) distance can be given. The center-to-center distance is the preferred alternative but a complementary indication is often necessary to avoid the confusion.

*F. On Electrochemistry*

The electrodes are made of conducting material, typically metal, but not necessarily. Pipette electrodes have been used very much in giant axons and for intra-cellular work. For practical reasons, metal electrodes are the rule for other applications. The problem with metal electrodes is that they carry electric currents as a flow of electrodes while electrons are hardly soluble in the hydrous solutions of the body. At the interface between both, an electrochemical process is necessary to transform the electric current in a displacement of ions instead of electrons. It is not possible to cover here the vast domain of electrochemistry but this has many consequences including the necessary biphasic stimulation pulses mentioned earlier. One important resulting issue for neural recording is that the electrode interface is often the major component of the electrode impedance and this impedance is not only directly responsible for the generation of input thermal noise but also for a reduction of the common mode reduction by introducing a variation factor in the electrode impedance values. The electrode impedance is also important for stimulation. A given level, of current density, the electrochemical changes at the interface become irreversible and the electrode will corrode which is not acceptable in chronic use. Also, if the electrode impedance is too high, it will take a significant fraction of the stimulator output voltage and limit the available stimulus intensity. Also variable electrode impedances introduce an uncontrolled variation in the stimulus intensity when the stimulator is voltage controlled. That is why a constant current source is preferred.

#### *G. Regarding electrode contact area*

As indicated above, the reversible stimulation current density is limited. This means that a larger electrode would allow passing larger currents. However, such larger currents are then distributed over larger areas and perhaps beyond the target, where they are wasted. The same is true for recording, where large contacts allow to reduce the electrode impedance but will average the signal over a larger area, including non-active regions.

The electrode material (often Platinum, or Platinum iridium in implants, silver/silver chloride for skin electrodes) has a major effect on the impedance as well as on the chronic biocompatibility of the electrodes. As a normal reaction to the implanted foreign material, a layer of fibrous tissue encapsulating the electrodes is systematically produced. This layer increases the distance between the electrode and the target tissue and adds a poorly conductive layer, all reducing the electrode sensitivity. It is not only the material that controls electrochemical processes and the foreign body reaction. Electrode surface treatments, the electrode micro-geometry and various coatings can play a major role.

#### *H. Regarding electrode impedances*

As a shortcut for techniques such as galvanometry, electrode impedance values are sometimes considered a reflecting the state of electrode interfaces. As a matter of fact, the current-tension relationship at the electrode interface is non-linear (current density and voltage dependent), frequency dependent, affected by hysteresis, dependent on the local chemical state among other issues. In other words, there is no unique value characterizing the electrode impedance. However, such a measurement is useful to check proper connections and as a kind of 'signature' of the setup being used.

Passing a very low alternating current to avoid damaging the recording electrode interface, in the frequency range of the signals to be recorded (or more typically, at 1 kHz) can provide a useful indication. For stimulation electrodes, measuring the potential reached while sending a stimulus pulse similar to the intended stimuli can do the same job. One should remember that the so-called impedance value obtained is for the full two-electrode loop, where the two electrode interfaces involved see the current with an opposite polarity and have thus a different contribution to the overall measurement.

#### *I. Regarding probes and leads*

Electrodes must be held in place and should make an electric contact with the tissue only over a selected area. Needles for example can be used as electrodes and will be insulated over their full length except the tip so that only the tip works as electrode area while the shaft is the lead carrying the electrical signal. The non-insulated diameter and length are necessary to calculate the contact area. A huge variety of devices have been developed to carry electrodes and maintain them in position. This goes from the familiar ECG skin surface gel-coated electrodes embedded in a piece of adhesive, to the various implantable cuffs maintaining a set of electrodes against the nerve around which they are implanted. Other electrodes penetrate the nervous system and the slanted electrode is a good example of an attempt to realize a 3D connection to individual fibers.

These devices are supposed to maintain mechanical stability of the contacts which is essential because micro-movements of the electrode have been shown to be responsible for activating inflammatory reactions. The lead necessary to carry away the electrical signal or bring in the stimulus can be responsible for tethering forces that must be avoided as well. This is why material softness is an important parameter as well. Even the currently used polymers do not seem to be soft enough for some applications.

All these aspects also touch upon the implantation technique where a good description of the electrode placement should be complemented with precise indications about fixation method. Accessibility can sometimes be a problem as illustrated by some attempts to develop intravascular electrodes.

#### *J. Electrode Biocompatibility*

Materials used to build the probes are usually well-known for their excellent bio-compatibility record. The mechanical properties seem often underestimated. On the other hand, it is important to remember that cleaning is an essential factor to avoid inflammatory reactions and is not to be confused with sterilization to avoid infections. Sterilization can affect the electrode surface structure. For all those reasons, cleaning and sterilization methods are worth mentioning if such issues are at stake.

### VII. The Recording Channel

In a data acquisition chain, recording is typically organized in a number of parallel channels with analog input converted to digital form and multiplexed into a data stream to a computer.

The input of each channel is typically a low noise high input impedance differential pre-amplifier of limited gain and low output impedance (e.g. [1]), followed by an RC high-pass filter [2]. This arrangement corresponds to the fact that the low amplitude electrophysiologic signals of interest are as a rule, superimposed on huge DC potentials such as the electrode interface potential [3].

The mains power frequency (50 or 60 Hz) is the most frequent and largest source of interference. Therefore, a hardware notch filter at that corresponding frequency can be used for some applications. Next, the signal Is amplified before to reach an anti-aliasing low-pass filter (cut-off frequency $F_c$ [4] that must eliminate all signal components at frequencies higher than half the sampling frequency ($F_s$). Thus $F_c<(F_s/2)$ [5]. For each time bin ($1/F_s$), the signal level is being held at a given constant level during the numeric conversion. Several channels run in parallel. The sampling can thus be simultaneous in all channels, waiting the end of all conversions before the next time bin is being sampled. Other systems sample channels successively introducing a short delay between channels.

The software of modern equipment can provide a large choice in numeric filtration algorithms (from IIR filters imitating their hardware counterparts to very sophisticated signal analysis techniques). The usage of the less optimal hardware filters should thus be limited to two circumstances: the absolute necessity to avoid signal saturation and the anti-aliasing phenomenon at A/D conversion. Most often, this can be done with the input RC and the anti-aliasing filter described above. However, additional hardware filters can be necessary in some circumstances. It should be remembered that physiological signals can be very difficult to distinguish from the overshoots generated by transient artifacts such as spikes or DC shifts high-pass filtered with a circuit of order>2 [6]. In addition, the use of single order high-pass filter allows to recover some information about DC shifts [7]. This is why EEG equipment use the RC value instead of a -3dB cut off frequency to characterize the high-pass filter, referred to as the 'time constant' while the -3dB cut-off is given for the low pass filter referred to as the 'filter'. This is good practice and should be maintained.

Note that amplifier characteristics such as precision and linearity are usually useless in neurophysiology because technical achievements at that level are much better that can be usefully exploited in the frame of physiology.

Input noise is a difficult topic because, even not considering the biological interferences, there are several technical sources of unavoidable noise with different characteristics. One should pay specific attention to the effect of input device current noise on the input circuit impedance, the thermal noise generated in the input circuit impedance, 1/F input noise and temperature input DC shifts.

### VIII. Supplemental Methods

To generate fig. 1B, an exponential growth relation was fit to the results of a Web of Science search for the search terms "TS=(brain (machine OR computer) interface) OR TS=((electrical OR optical OR acoustic) neuromodulation) OR TS=(neural interface)" for all item types between the years 1990 and 2019. This search returned 27,188 records, including 13 items prior to 1990. The literature for nerve stimulators (search term "TS=(Nerve stimulation)") is much larger (70,473 records) but has been growing more slowly, doubling every 8.6 years since 1960.
